# Morphological variability in the mucosal attachment site of *Trichuris muris* revealed by X-ray microcomputed tomography

**DOI:** 10.1101/2021.01.07.425704

**Authors:** James D. B. O’Sullivan, Sheena M. Cruickshank, Philip J. Withers, Kathryn J. Else

## Abstract

Parasitic infections can be challenging to study, because light and electron microscopy are often limited in visualising complex and inaccessible attachment sites. Exemplifying this, *Trichuris* spp. inhabits a tunnel of epithelial cells within the host caecum and colon. A significant global burden of this infection persists partly because available anthelminthics lack efficacy, although the mechanisms underlying this remain unknown. Consequently, there is a need to pioneer new approaches to better characterize the parasite niche within the host and investigate how variation in its morphology and integrity may contribute to resistance to therapeutic intervention. To address these aims, we exploited 3D X-ray micro-computed tomography (microCT) to image the mouse whipworm *T. muris* in caeca of wild-type C57BL/6 and SCID mice *ex vivo*. Using osmium tetroxide staining to effectively enhance contrast of worms, we found that a subset exhibited preferential positioning towards the bases of the intestinal crypts. Moreover, in one rare event, we demonstrate whipworm traversal of the lamina propria. This morphological variability contradicts widely accepted conclusions from conventional microscopy of the parasite niche, showing *Trichuris* in close contact with the host proliferative and immune compartments that may facilitate immunomodulation. Furthermore, by using a skeletonization-based approach we demonstrate considerable variation in tunnel length and integrity which may represent an indicator of tunnel “health”. The qualitative and quantitative observations provide a new morphological point of reference for future *in vitro* study of *Trichuris-*host interactions and highlight the potential of microCT to more accurately characterise enigmatic host-parasite interactions.

**Author Summary:** Parasites are often difficult to observe once established within host tissues, presenting a barrier to biological understanding and therapeutic innovation. Whipworms (*Trichuris* spp.) affect 500 million people worldwide, causing significant disability, and appear partially resistant to widely used “deworming” drugs. However, the inaccessibility of worms within the cells of the host intestine makes them highly challenging to image and study. By investigating *Trichuris* attachment sites in 3D, using X-ray micro-computed tomography, we found that the niche is highly variable in size and, contrary to reports in all previous studies, can also penetrate different layers of intestinal tissue. By showing that worms are positioned much closer to host immune cells that previously appreciated, we provide a morphological reference point for future studies on how *Trichuris* effectively avoids clearance by the host. The non-invasive imaging approach used represents an excellent opportunity to clarify the lifecycles of other difficult-to-study parasites.

## 1. Introduction

Microscopy and histology are key tools in understanding the life cycles of parasites. However, when parasite life cycles involve establishment within host tissues and even cells, identifying and characterizing regions of interest using 2D sectioning becomes challenging. Exemplifying this is *Trichuris* spp., the causative agent of trichuriasis, a neglected tropical disease estimated to affect almost half a billion people worldwide as of 2010 [1]. *Trichuris* spp. has a unique life-cycle in which the adult worm lives with its anterior two thirds embedded within an “intracellular” tunnel within the columnar epithelium of the host caecum [2,3]. This embedded portion of the worm houses the bacillary band and the stichosome, both of which are specialized organs thought to facilitate this intracellular lifestyle [4,5]. Like many other endoparasites, the mechanisms underlying niche formation and maintenance are poorly defined, due to the challenging nature of examining a complex 3D interface with the host using only 2D sections characteristic of conventional histology and electron microscopy. Even if worms are successfully located within the tissue, a non-ideal slice orientation can preclude informative imaging. 3D imaging approaches provide an opportunity to overcome these limitations and assist in understanding the attachment site *Trichuris*, as well as other clinically relevant parasites. However, observation of *Trichuris* within the niche using confocal, multiphoton or light sheet microscopy is severely limited due to the genetic intractability of the parasite and challenging structure of the intestine [6], so other approaches are required.

Existing anthelminthic drugs lack efficacy in controlling *Trichuris* infection [7–9]. Therefore, continuing development of novel anthelminthics remains an important contemporary objective in *Trichuris* research [10]. *Trichuris muris*, which infects mice, is commonly used as a model to investigate trichurid host-parasite interactions and therapeutic strategies due to its well-characterized development and the genetic tractability of its host [11,12]. Preclinical drug and vaccine trials have consistently shown partial reduction of *T. muris* worm burdens [13–15], suggesting that worms within the same host are differentially susceptible to their delivery, toxicity or immunogenic actions. Why some worms are more susceptible to therapeutic intervention than others remains unclear, but the phenomenon may be related to the success of niche maintenance, including the integrity of the host epithelium overlying the worm. Improving our understanding of the *Trichuris* survival within the host, and examining how the niche varies between worms during chronic infection is therefore an important objective for therapeutically addressing worm resistance [11,12].

Most of the knowledge of the trichurid tunnel structure and composition so far has been obtained by electron and light microscopy. Adult worms appear to occupy a tunnel of ‘dead’ epithelial cells, with bacteria and damaged, fragmented organelles present in the more posterior tunnel regions [2,5]. The survival of *Trichuris* within this niche in thought to be promoted by modulation of the host immune response. Specifically, secreted proteins [16] promote a pro-inflammatory milieu whilst suppressing elevated epithelial turnover, a key immune effector mechanism which ejects *Trichuris* from its niche [17,18]. Despite an increasingly complete picture of the immunologic bases for host susceptibility and resistance [12], the nature of tunnel formation and development remains mysterious. During the earliest hours of invasion, larvae actively interact with the live cells at the base of the crypts of Lieberkühn to form the multi-intracellular niche [19,20]. After the sequential moults leading to adulthood, the tunnel becomes much longer and more tortuous, presenting an obstacle to effective imaging [3]. However, microscopic studies of niche morphology to date have all concurred that adult *Trichuris* live exclusively within the epithelial cells at the surface of the mucosa, directly adjacent to the lumen [2,3,21,22]. X-ray micro-computed tomography (microCT), which is inherently 3D and is capable of providing images of a comparable resolution to histological approaches [23,24], provides unique opportunities to interrogate the structure and morphology of the tunnel in detail [25]. Capturing a volume, rather than sampling a random 2D plane, offers the opportunity to accurately characterize the morphology of multiple attachment sites in their entirety, as well as the degree of variation between them, potentially revealing new information about the *Trichuris* life cycle.

By exploiting microCT imaging at “histological” resolutions, we visualized tunnel morphology in the developing and adult worms. The aims of the study were to 1) describe the 3D morphology of the attachment site throughout worm development; and 2) to quantify the degree of morphological heterogeneity across multiple *Trichuris* attachment sites during chronic infection. By exploiting the 3D nature of the data, we morphometrically explored multiple tunnels using a skeletonization-based method order to estimate tunnel length, which we suggest may constitute a basis for future *in situ* screening tools of anthelminthic drugs. We also describe the presence and frequencies distinct sets of tunnel morphologies. To achieve these goals, we developed a sample preparation protocol adopted from scanning electron microscopy which facilitates quantitative analyses, is well-suited for the study of the *Trichuris* attachment site and lends itself to analyses of other hard to image parasites within their hosts

## 2. Results

### 2.1. Morphology of the worm and host attachment site in MicroCT

In order to investigate qualitatively the morphology of the epithelial tunnel throughout the development of chronic infection in the caecum, we examined the caeca of mice at different stages of infection, roughly corresponding with the *T. muris* moulting pattern. Male C57BL/6 mice were infected orally with a low dose of *T. muris* eggs in order to ensure a chronic infection [26]. At 7, 14, 21 and 35 days post-infection (PI) with *T. muris*, 3-5 mm tubular fragments of caecum were dissected and processed for microCT imaging (**Fig. 1 A**). In caecum fragments taken 35 days PI, we identified an optimal staining protocol utilizing Osmium Tetroxide (OsO_4_), which allowed superior contrast between the worm and surrounding tissue when compared to worms immersed in aqueous potassium triiodide (Lugol’s Iodine) and PTA (**Fig. 2 A-C; Table 1**).

**Table 1:** Contrast enhancement approaches for microCT of T. muris-containing caeca.

| Stain | Composition | Mounting |
| --- | --- | --- |
| $\text{I}_2\text{KI}$ (Lugol's iodine) | 0.6% potassium iodide,<br>0.3% iodine in distilled<br>water | suspended in distilled water |
| PTA<br>(Phosphotungstic<br>Acid) | 1% in 100% ethanol | suspended in 100% ethanol |
| $\text{OsO}_4$ (Osmium<br>Tetroxide) | 1% in distilled water | Dehydrated<br>Critical point dried |

**Figure 1.**
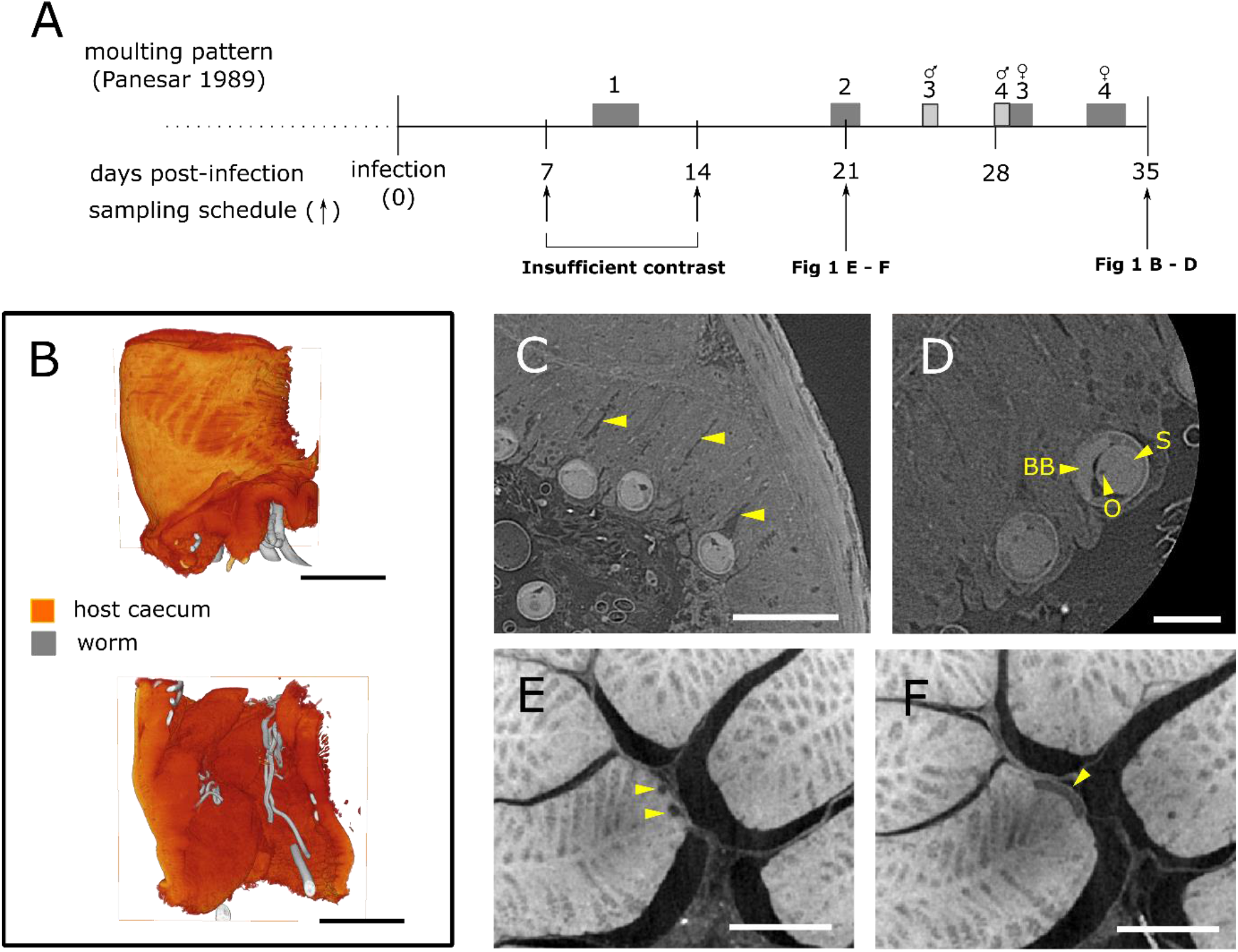
Visualising parasite and host gut morphology in 2D slices and 3D renderings during the development of chronic trichuriasis. Fragments of caecum were dissected from male C57BL/6 mice at days 7, 14, 21 and 35 post-infection with <40 T. muris eggs. The caecum was OsO_4_-stained and critical point dried before being mounted and imaged with microCT. **A**) Timeline showing the moulting pattern of Trichuris muris and the sampling points for the caeca used in this study **B**) Volume renderings (full and half-cutaway) showing positioning of worms within the caecal biopsies. **C**) Slice of day 35 PI caecum showing locations of Crypts of Lieberkühn (yellow arrowheads). **D**) High resolution slice of day 35 PI caecum showing the oesophagus (**O**), bacillary band (**BB**) and stichocyte (**S**) of an adult worm. **E**) slice of a day 21 PI caecum fragment showing two parts of a single L3 worm (yellow arrows), negatively contrasted against the host mucosa. **F**) slice showing positioning of the single L3 worm (yellow arrow) at the surface of the host mucosa. Scalebars B = 300 µm, C = 100 µm, D, E, F = 500 µm.

**Fig 2:**
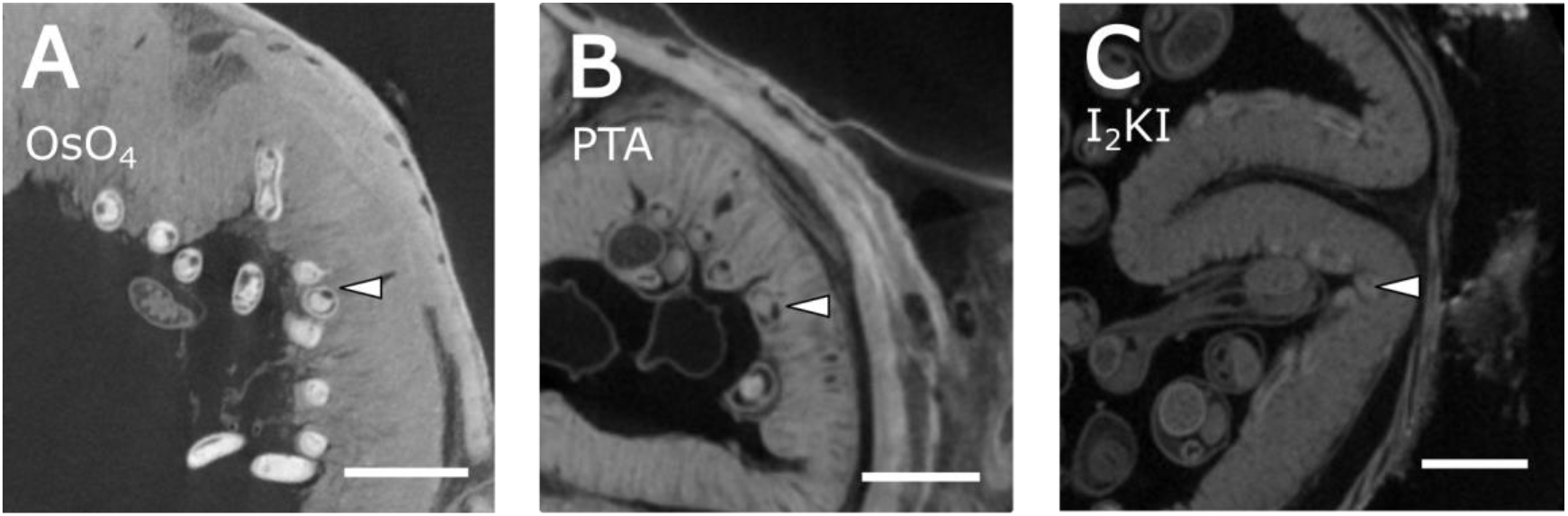
Staining with Osmium tetroxide is more effective than either Phsophotungstic Acid (PTA) or Lugol’s Iodine (I_2_KI) in generating contrast of the stichosome in adult (day 35 post-infection) worms. Samples were **A)** Stained in osmium tetroxide and critical point dried **B**) Stained in PTA and embedded in 4% gelatin or **C**) Stained in Lugol’s iodine and embedded in 4% gelatin before imaging. Worms (white arrowheads) appeared better contrasted in OsO_4_-stained caeca, perhaps due to the highly membranous nature of the stichosome. Histogram minima were all assigned to the material surrounding the caeca. Scale bars: 100 µm.

In caeca from C57BL/6 mice taken 35 days PI, microCT scanning of whole fragments (pixel size = 3.62 µm, **Fig. 1 B**) revealed the attachment sites of multiple worms. When the magnification was increased in order to visualize regions of interest in more detail (pixel sizes 1.56 µm and 0.59 µm), multiple aspects of worm and host morphology were apparent. The crypts of the host mucosa were visible as layers of reduced X-ray attenuation (**Fig. 1 C**). The bacillary band, stichocytes and oesophagus of the worm and their orientations were all distinct, exhibiting higher X-ray attenuation than the surrounding host tissue (**Fig. 1 C, D**). Cuticular inflations, another specialized anterior organ [27], were not distinguishable in the adult worms, either in slices or volume rendering of the worm and attachment site. Larval worms at days 7 and 14 post-infection were not observed in tissue samples taken from these time points, we speculate due to lack of contrast. However, one worm was observed in the tissue sample taken 21 days post-infection (**Fig. 1 E-F**). In contrast to the day 35 PI adult worms, the day 21 PI L3 larvae were negatively contrasted against the host mucosa, perhaps due to the absence of a well-developed stichosome.

### 2.2. Worms exhibit “head down” behavior

In the lengths of caecum dissected from C57BL/6 mice, we identified variation in the positioning of adult worms (day 35 PI) within the niche. The majority of the worms observed (6/10; 60%) were positioned with the anterior-most portion of the stichosome pointed down the crypts of Lieberkühn, with the tip of the head close to the muscularis mucosae (**Fig. 3A - D**), a minority (3/10; 30%) were positioned in the conventionally understood manner with the tip of the head lying next to the lumen in the epithelial cells at the apices of the crypts (**Fig. 3I - L**). In addition, one worm was observed with a “curled” head morphology, where the anterior-most end of the worm was positioned within the crypt, with the tip of the head curled back towards the crypt apex (**Fig. 3E - H**). In a separate mouse host, the positioning of the tip of the head near the proliferative compartment during “head down” behavior was verified by light microscopy using BrDU (5-bromo-2’-deoxyuridine) immunohistochemistry to label actively proliferating cells, as well as with Haematoxylin and Eosin paraffin histology. However, the depth to which the tip of the head reaches is less evident than in light microscopy than microCT images due to the orientation of the slicing plane relative to the tissue (**Fig. 4**).

**Figure 3:**
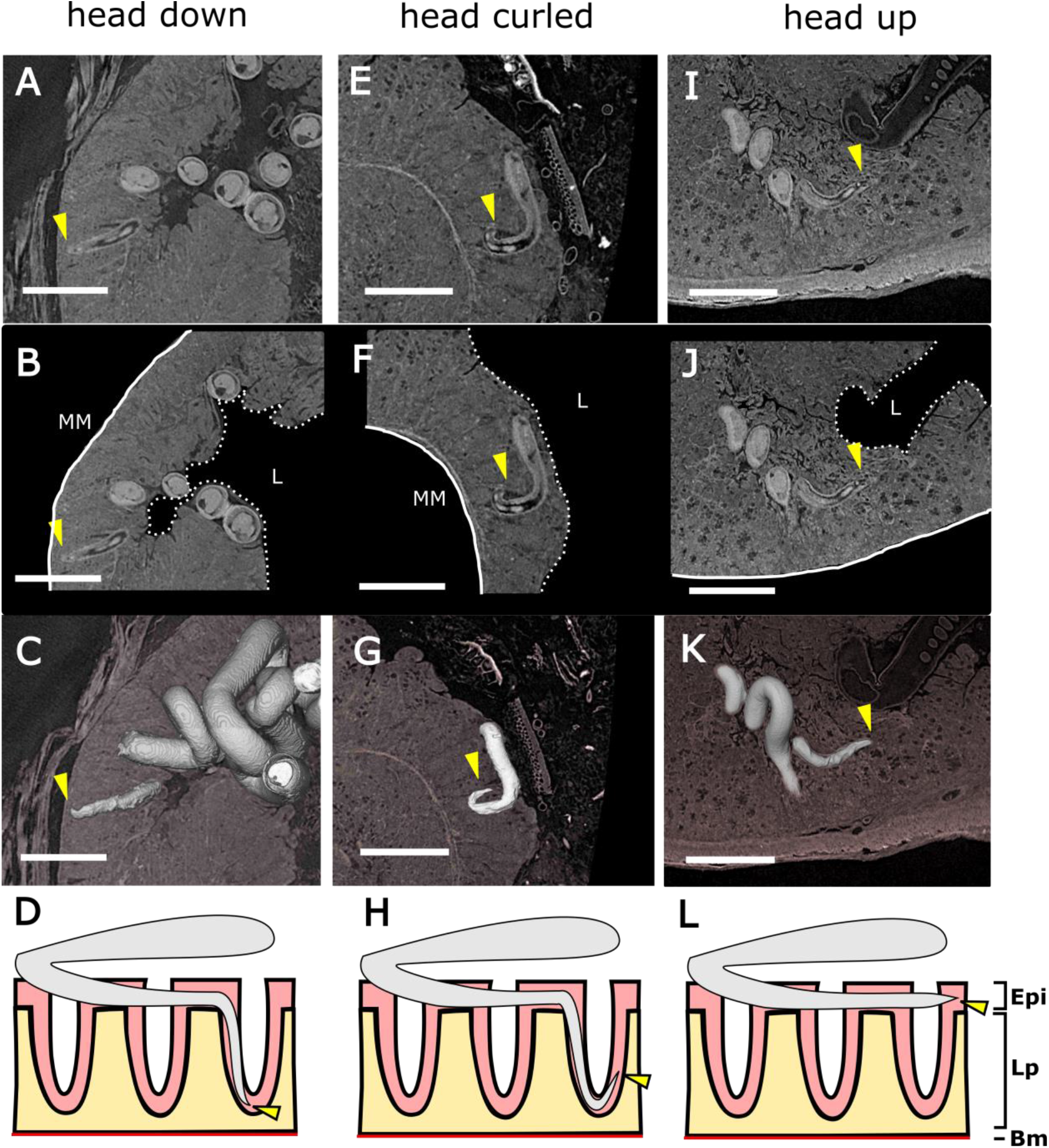
Variable positioning of the anterior stichosome at day 35 post-infection in a C57BL/6 mouse. Representative images of the OsO4-stained caecum from mice at day 35 post-infection with a low dose of Trichuris muris showing the different patterns of head behavior. Worm positioning was visualised and is depicted here in slices, volume rendering and diagrammatic format. Three sets of morphologies were revealed among the ten worms investigated. Some worms demonstrated a “head down” behavior, in which the head is pointed down the crypt towards the muscularis mucosae. This behavior is shown in a slice (A), with the tip of the head indicated by a yellow arrow and in (B) is shown in relation to the muscularis mucosae (MM) and lumen (L). C) Shows a volume rendering of the worm stichosome (grey) in relation to the host mucosa (pink). D) Illustrates the positioning of the worm within the attachment site (Epi – Epithelium; Lp – Lamina Propria; Bm – Basement Membrane). One worm within the sample exhibited a “head curled morphology”, in which the head region went down the crypt, and then turned back on itself towards the lumen, shown in slice (E) schematic (F) volume rendering (G), and illustrated (H) views. In the “head up” morphology, shown in slice (I) schematic (J) volume rendering (K) and illustrated (L) views, worms were positioned as conventionally understood, with the tip of the head within the epithelial cells directly adjacent to the lumen. All scale bars = 600 µm.

**Fig 4:**
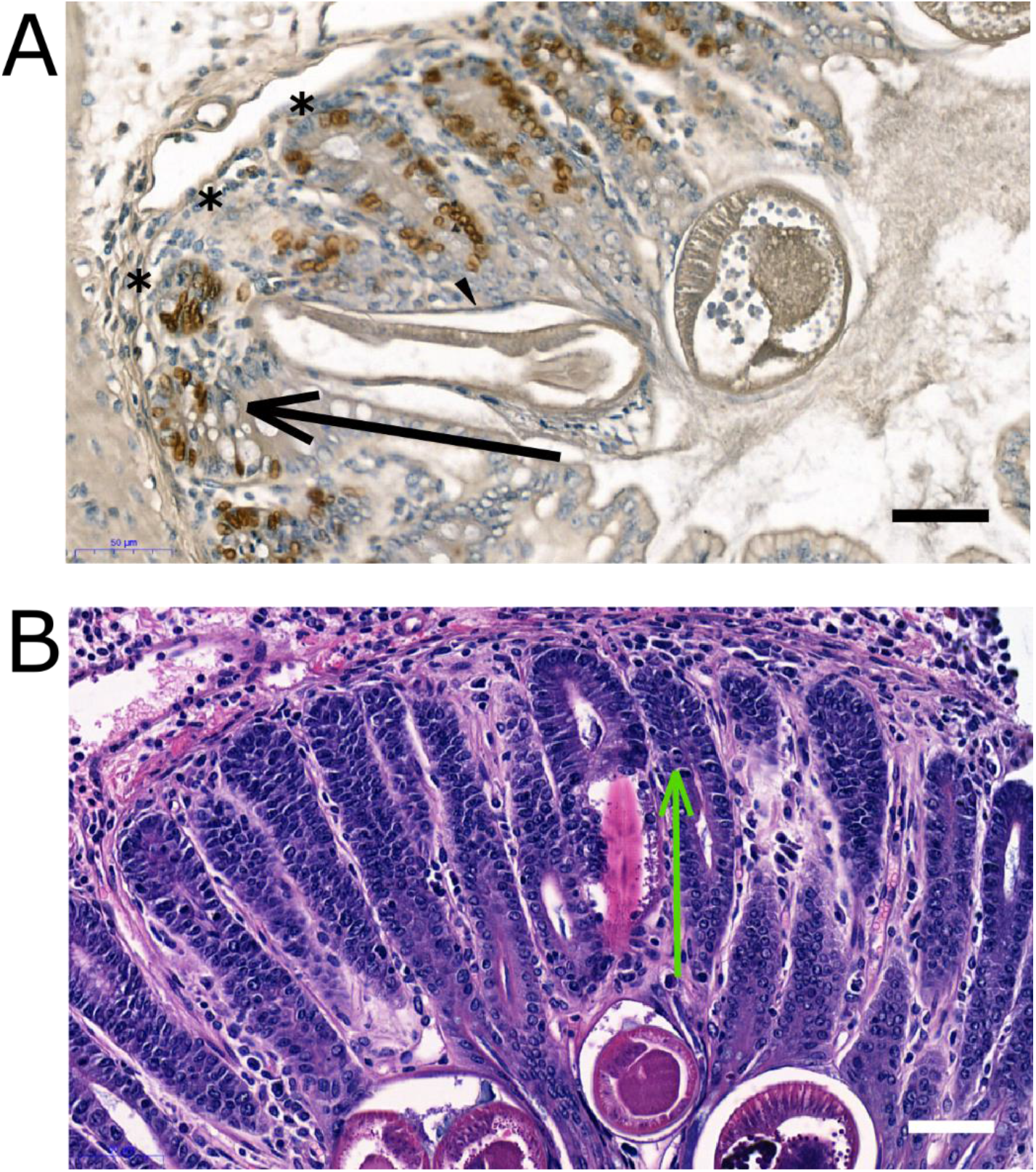
Histological observation of head-down behavior. Formalin-fixed paraffin-embedded tissue was exhaustively searched for evidence of head-down behavior, yielding two examples **A**) BrDU immunohistochemistry shows the proximity of the worm anterior (orientated in the direction of the black arrow) relative to the proliferative cells (brown) at the base of the epithelial crypts (*). The worm here appears to be crossing the lamina propria (black arrowhead). **B**) Haematoxylin and Eosin staining again showing the orientation of the worm anterior (green arrow, eosinophilic cuticle visible) towards the base of the crypts. Here the worm appears confined to the epithelial cell layer. Scale bars = 50 µm.

**Figure 4:**
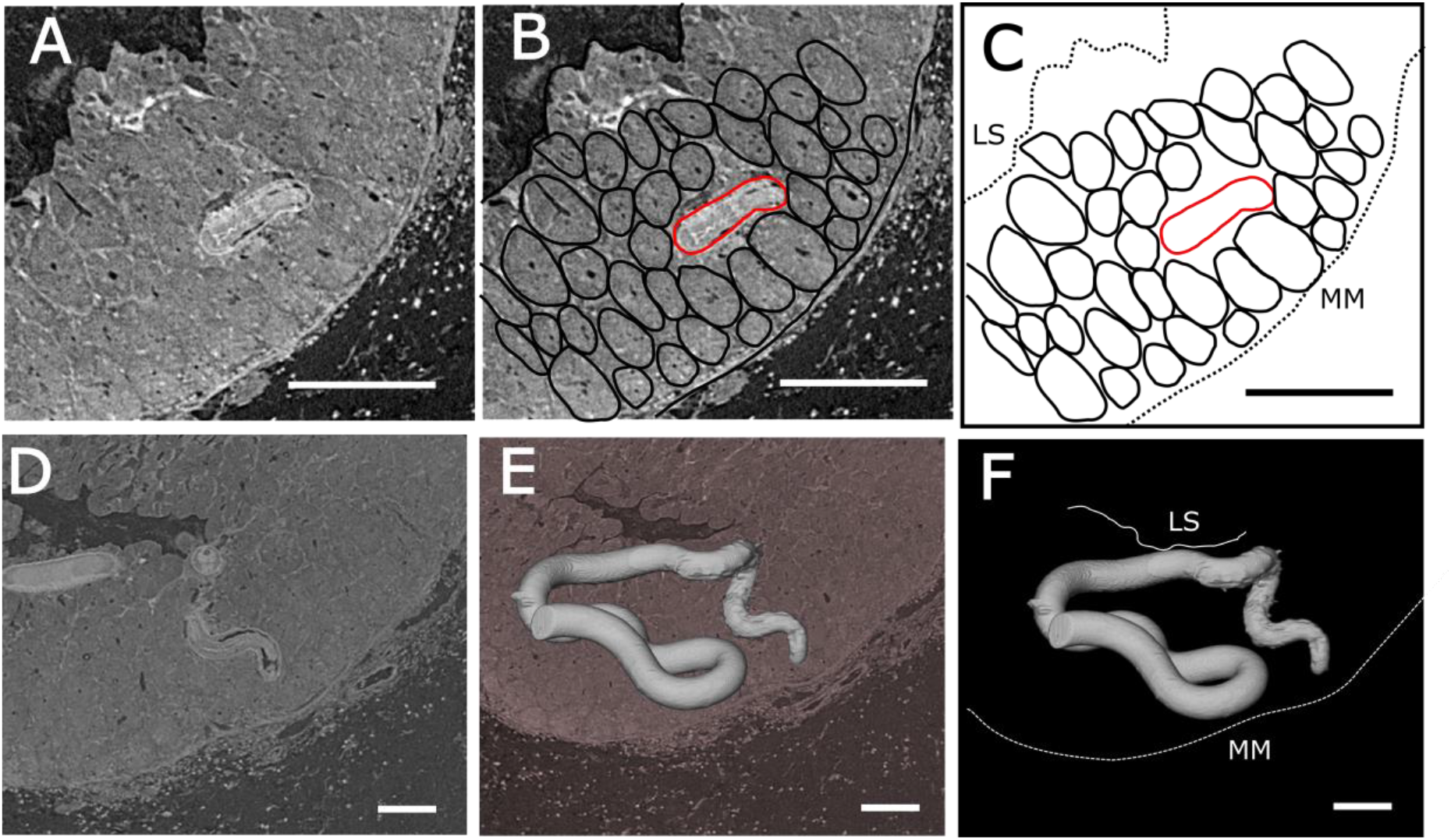
Worm head morphology in a SCID mouse reveals a worm traversing a crypt at 35 days post-infection. A SCID mouse caecum dissected at day 35 post-infection and prepared using OsO_4_ staining and critical point drying. The caecum contained one worm exhibiting highly unusual positioning. **A**) The worm is travelling laterally across the mucosa, over a distance roughly equivalent to three crypts. **B**) The worm is highlighted in red and the positioning of the crypts is represented by black ovals. **C**) The positioning of the worm (red) is shown in schematic form relative to the lumenal surface of the mucosa (LS) and the muscularis mucosae (MM). **D**) Slice showing the zig-zagging positioning of the worm head. **E**) Volume rendering of the worm showing the positioning of the whole stichosome region compared to the surrounding mucosa (pink, slice). **F**) Schematic showing the spatial relationship between the worm stichosome (grey), the luminal surface of the mucosa (LS) and the muscularis mucosae (MM). Scale bars = 150 µm.

In addition to C57BL/6 mice, we were interested in observing the positioning of worms in Severe Combined Immunodeficient (SCID) mice. These mice lack the adaptive components of the immune response, and are susceptible to *T. muris* at both high and low doses. In this study, SCID mice were infected with a low dose (<40 eggs) of *T. muris* in order to replicate the infection burden of the C57BL/6 mice examined in **Figs 1 + 2**. When microCT of SCID caeca was carried out using Osmium tetroxide staining, further head-down morphology was observed, similarly to C57BL/6 mice. However, in one SCID mouse we observed one worm which appeared to travel laterally between crypts, half-way between the lumenal surface and the muscularis mucosae (**Movie 1**). Closer inspection under higher magnification (pixel size = 1.16 µm) confirmed this lateral movement of the worm through the crypt including penetration of the lamina propria (**Fig. 4A-F**).

In order to examine the occurrence of the head-down behavior across mouse strains and sexes, we undertook a series of scans using the staining methodology described above. Frequencies of head down behavior in C57BL/6 (male, female) and SCID (female only) mice are recorded in **Table 2**. Statistical analysis was not attempted due to the low frequencies of worms sampled in each group. However, in the 13 mouse donors examined, we note that qualitatively there was no trend evident in the occurrence of the head down behavior between sexes or strains.

**Table 2:**
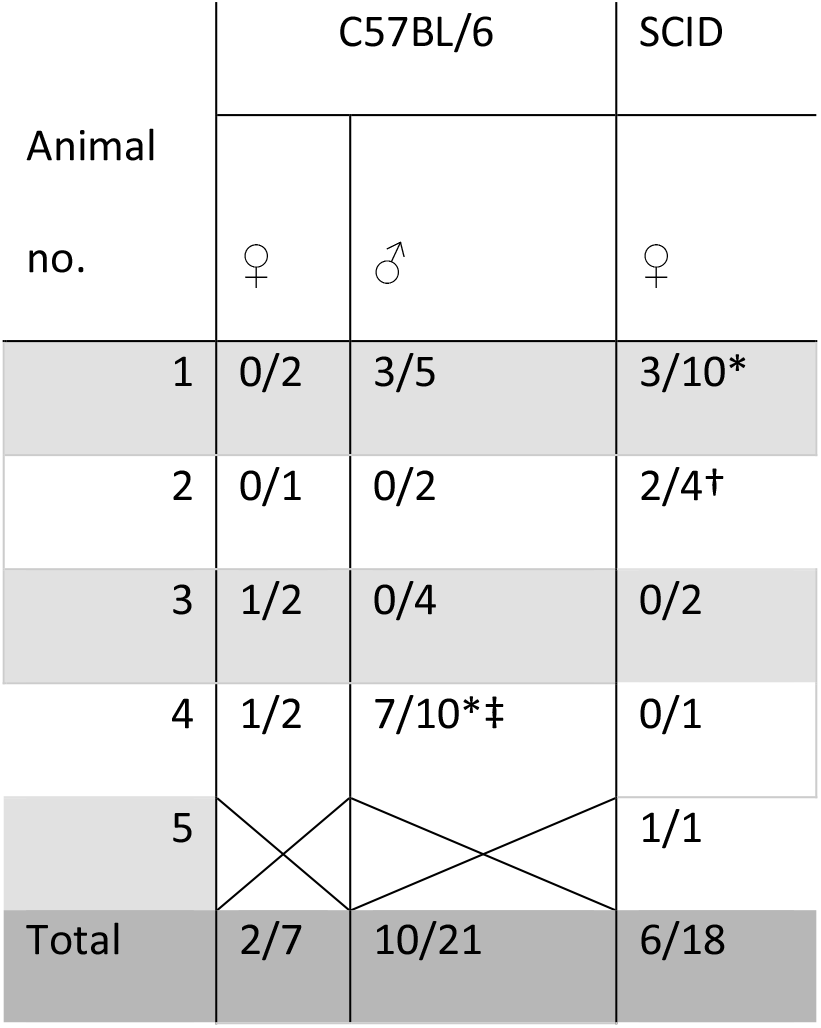
Proportions of head down behavior observed in C57BL/6 and SCID mice (each cell is a separate individual; total = 13 mice), expressed as number of head down worms observed/total worms observed per animal. In cases marked with *, multiple segments (3 each) of caecum were observed and pooled. ‡ indicates the C57BL/6 mouse from which observations were recorded in **Figures 1, 3** and **5**. † indicates the SCID mouse from which the unusual head positioning in **Figure 4** was recorded.

| Animal<br>no. | C57BL/6 |  | SCID |
| --- | --- | --- | --- |
|  | ♀ | ♂ | ♀ |
| 1 | 0/2 | 3/5 | 3/10* |
| 2 | 0/1 | 0/2 | 2/4† |
| 3 | 1/2 | 0/4 | 0/2 |
| 4 | 1/2 | 7/10*‡ | 0/1 |
| 5 |  |  | 1/1 |
| Total | 2/7 | 10/21 | 6/18 |

### 2.3. Skeletonisation analysis permits the *in situ* length of the epithelial tunnel to be quantified

In order to understand whether heterogeneity in the niche could also be extended to include variations in the integrity and lengths of the tunnels themselves, we utilized skeletonization to quantitatively assess the number and size of previously described “breaks” in the epithelium covering the tunnel [3], as well as the length of the embedded portion of the stichosome. In the caecum images from the C57BL/6 mouse at day 35 P.I. examined in **Fig 1** and **Fig 3**, worms were assigned a unique ID and subsequently the proportions of the stichosome which were covered and uncovered by the epithelial layer measured. Worms were segmented, and skeletonization facilitated length measurements of the tortuous stichosome (**Fig. 5 A-C**). The skeletonized worm was manually fragmented based into “covered” and “uncovered” regions.

**Figure 5:**
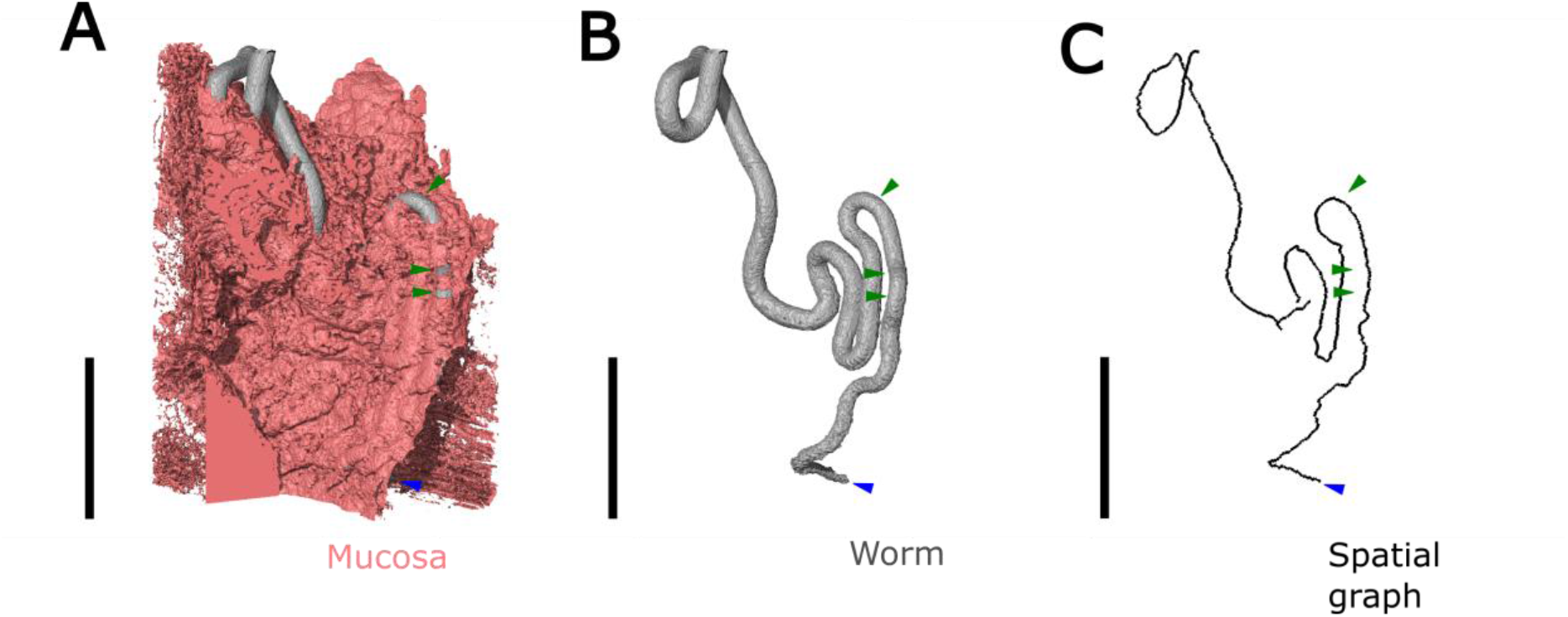
Skeletonisation approach for measuring the epithelial coverage of the syncytial tunnel of T. muris. Representative images taken of the caecum of one mouse (n = 7 worms were measured) infected with a low dose of T. muris, at day 35 PI. The caecum fragment was prepared for microCT by staining with OsO_4_ and critical point drying. After imaging, the worms and the surrounding mucosa were segmented and skeletonisation was used to analyse the integrity of the epithelial tunnel. **A**) Image showing a worm (grey) and the epithelium (pink) covering the tunnel. Green arrows indicate breaks in the epithelium covering the tunnel, which leave the worm exposed to the lumen. The blue arrow indicates the position of the worm head. **B**) Surface rendering showing a worm in isolation (grey). **C**) Rendering of the skeletonised worm “spatial graph” which was used to estimate the lengths of worms left exposed by breaks in the tunnel covering. All scale bars = 500 µm.

Therefore, the length of worm covered by the epithelial tunnel could be quantified, and was expressed as a percentage as “epithelial coverage”. Epithelial coverage was measured in seven worms in the caecum of the male C57BL6 mouse (pixel size = 3.619 µm), and was highly variable; of the seven worms analyzed only one had its stichosome totally covered by epithelium, with the other six worms having between 3. 07% and 17.44% of the stichosome exposed (**Table 3**).

**Table 3:** Estimated tunnel lengths for seven worms at d35 PI in a chronically infected C57BL/6 mouse. The lengths of breaks in the epithelium, and thus the proportions of tunnels which were exposed were estimated by using AVIZO’s centreline tree algorithm to measure segmented worms.

| Worm ID | total tunnelled length including epithelial breaks (mm) | cumulative length of exposed stichosome (mm) | Number of breaks in epithelium | cumulative length of unexposed stichosome (mm) | proportion of stichosome exposed (%) |
| --- | --- | --- | --- | --- | --- |
| 4-roi2 | 3.13 | 0.18 | 2 | 2.96 | 5.82 |
| 6-roi3 | 7.76 | 0.00 | 0 | 7.76 | 0.00 |
| 9-roi5 | 3.99 | 0.70 | 4 | 3.30 | 17.44 |
| 12-roi6 | 7.41 | 0.78 | 3 | 6.63 | 10.52 |
| 13-roi7 | 7.49 | 0.95 | 2 | 6.53 | 12.73 |
| 14-roi7 | 9.25 | 0.28 | 2 | 8.96 | 3.07 |
| 15-roi7 | 7.41 | 0.94 | 5 | 6.47 | 12.68 |

The cumulative tunneled length of the stichosome calculated for seven worms (**Table 3**), was also highly variable, ranging from 2.96 mm to 8.95 mm. Furthermore, the number of breaks varied between worms (range: 0 to 5). Importantly, the tunneled length of the stichosome was not always representative of the total length of the stichosome. In all worms, some length of the posterior-most region of the stichosome was positioned outside of the epithelial tunnel within the intestinal lumen. Often the stichosome was not fully encompassed within the imaging field of view, preventing measurement of the total stichosome length. However, at day 35 post-infection, previous work has shown the lengths of *T. muris* to range between 12 – 16 mm [28]. Given that the trichurid stichosome was accounts for two thirds of the total body length [29], we therefore demonstrate that the proportion of tunnel stichosome buried within the host relative to that hanging free in the lumen ranges widely between worms.

## 3. Discussion

Emerging 3D imaging techniques including microCT offer opportunities to better understand the nature of attachment sites within intact tissues and bypass the unreliable sampling of features inherent in 2D slicing approaches. Here we have presented a case study, using the parasitic intestinal nematode *Trichruis muris*, which demonstrates how thorough exploration of the host parasite interface in 3D can enhance understanding of parasite life cycles. We report two discoveries; firstly, that there is significant variation in the “shape” of the attachment site, because a subset of worms appear to preferentially position their heads toward the proliferative compartment of the crypts and even invade the lamina propria. Secondly, we show through 3D quantification that there is also significant heterogeneity in tunnel length and integrity of the overlying epithelial tissue between worms.

Previous attempts using histology, TEM and SEM have provided a partial view of the intra-epithelial tunnel, both from cross-sections and the luminal surface [2–5,27]. However, 3D imaging using microCT has allowed us to catalogue a more diverse set of sub-surface morphologies than previously appreciated, with implications for our understanding and future study of host-parasite interactions underlying the immunopathology of chronic trichuriasis. Whilst microCT has previously been used to qualitatively assess the morphology of endoparasites within soft tissues [30], here we carried out a thorough, semi-quantitative exploration of the parasite attachment site over multiple hosts. In order to produce effective imaging results, osmium tetroxide staining proved particularly suitable in contrasting adult worms against the host epithelium. We speculate that this is due to the membrane-rich and therefore highly osmophilic nature of the stichosome ultrastructure, which has been noted previously by our lab via 3D electron microscopy [31]. Given the growing interest in using microCT for imaging biological soft tissues [32], and the need for the development of effective soft tissue imaging protocols, we have demonstrated OsO_4_ as a useful and specific stain for *Trichuris*, which may even be extended to the CT visualization of other highly membranous cellular structures such as acinar and beta cells of the pancreas and plasma cells [33–35].

The “head-down” and “head curled” behaviors have not been previously described in *Trichuris* spp., and provide new context to existing understanding of the immunomodulatory actions of the worm. Notably, these results contradict the widely accepted conclusions that adult *Trichuris* only lives within the most apical epithelial cells adjacent to the lumen [2,3,20,22], and highlight that caution should be adopted in interpreting pioneering histological studies of endoparasites, given the disadvantages of sampling complex attachment sites in 2D. The observations here have provided a clear context for future *in vitro* and *in vivo* work. Namely, studies that aim to pry apart the basis of immunity and establishment of chronic infection should now consider that *Trichuris* may be in direct contact with both the lamina propria and intestinal stem cells. For example, in previous discussion of worm modulation of epithelial hyperproliferation, it was reasoned that relevant immunoregulatory compounds must be secreted, because the worms do not come into contact with the proliferative epithelial cells at the base of the crypt [36]. However, here we show conclusively that worms exhibiting head-down behavior come into close contact with, and probably directly damage, proliferating cells labelled with BrDU. We have shown that head-down behavior occurs in two different host immune environments, both including (C57BL/6) and lacking (SCID) the adaptive arm of the immune response critical to worm expulsion. Regarding the functional significance of the behavior, we speculate that the head-down behavior may suppress proliferation of the epithelial stem cells at the crypt base in a manner conducive to niche maintenance. Alternatively the head-down and head-curled behavior may be two stages of a single process, by which worms seek nutrition and extend their tunnels by appropriating the cell cytoplasm at the crypt base, rather than by invading only the cells adjacent to the lumen as previously thought [3].

In this work, we demonstrated the utility of microCT in identifying rare events by screening multiple tissue samples. In one such event (identified only once in microCT) occurring in the caecum of a SCID mouse, we observed traversal of adjacent crypts by the worm, showing direct contact between the worm and the lamina propria. The extreme rarity of this event, even with the benefits of imaging many tissue fragments in 3D, would render it virtually unobservable by sectioning-based histology. This behavior is presumably facilitated by the action of the sharp worm stylet and digestive proteases [37]. In the pathology of high-burden trichuriasis, anaemia and damage to the host gut is seen [38] and thus traversal of the lamina propria could account for severe pathology. Perhaps breaching of the intestinal barrier promotes a pro-inflammatory response conducive to worm survival [12]. A pertinent outstanding question is whether both crypt traversal and head down behavior occur in the much thicker human mucosa during trichuriasis (*Trichuris trichiura*). The absence of these behaviors would represent a significant divergence in the life cycle between the murine and human parasites, while their presence may be responsible for the pathology of heavy chronic infection.

In addition to the morphology described here, we also developed a method by which the tunneled length of the worm may be measured in 3D. The process of expulsion which occurs upon successful resolution of infection is not yet possible to visualize using the *ex vivo* approaches currently used in screening for anthelminthic efficacy. We believe that measurement of tunneled length is a useful proof of concept for future research into therapies for trichuriasis, in particular as a measure of resistance or therapeutics; we may hypothesize that the shorter the tunneled length, the more the processes of tunnel formation and maintenance are compromised by the relevant anthelminthic. Indeed, it may be that those worms which we observed with a short tunneled proportion of stichosome may be in the process of expulsion. Similar to a previous study by SEM [3] we observed breaks in the epithelium covering almost all of the worms analyzed. These are also prospective measures of susceptibility to anthelminthic, since they represent the extent to which the worm’s metabolically active stichosome is directly exposed to anthelminthics in the intestinal lumen. SEM has been an important tool in assessing drug efficacy *in vitro* [39], and a recent development of drug screening utilizes high-throughput quantification of worm motility *ex vivo* [40,41]. However, current methods lack any assessment of the status of the worm niche *in situ*. 3D imaging of the parasite can potentially fill this gap and increase the sensitivity of drug screening approaches [10,25], for example by correlating *ex vivo* worm motility with the average length of tunnels *in situ*.

In a broader context, we have demonstrated how microCT offers an opportunity to reassess the life-cycles of other parasites, and characterize intra-tissue migration and establishment with additional clarity. For example, *Ascaris* is another soil transmitted helminth which migrates through the lungs and, and 3D imaging of the distribution of worm larvae may clarify preferential routes in this poorly-understood migration [42]. Likewise, flatworms of the genus *Schistosoma* migrate through skin and lungs, with chronic infection featuring egg-associated granulomas in the liver, intestine or bladder and other organs [43]. Again, the 3D distribution of eggs and migrating larvae, as well as remodeling of host tissue i.e. angiogenesis may be particularly susceptible to investigation by microCT [25]. Therefore, microCT offers an opportunity to revisit classic parasitological discoveries in a modern context, in order to address challenging remaining questions in the study of host-parasite interactions. More broadly, the capabilities discussed may be applicable across a great variety of cases where heterogeneity of tissue during disease obfuscates accurate characterization of biopsies by conventional means [44].

## 4. Conclusion

Here we have used microCT, to provide a novel appreciation of *T. muris* positioning within its attachment site. By exploiting 3D visualization of the worm within its “epithelial” tunnel using effective sample staining, we have revealed qualitatively and quantitatively that the attachment site is heterogeneous in size and morphology, and that worms appear to contact directly the epithelial stem cells and lamina propria. Beyond *Trichuris*, these findings highlight there may be other similarly surprising morphological features in other nematode or flatworm parasites which inhabit animal tissues; indeed, further investigation of such features using 3D approaches may be a rich source of future biological insights and therapeutic opportunities that address the continuing burden of NTDs. Perhaps, in the footsteps of those who discovered and described these fascinating pathogens, it is time to revisit and update our understanding of parasite morphology using newly available techniques, in order to best contextualize our recent progress in understanding the immunology of these infections.

## 5. Materials and Methods

### 5.1. Experimental Animals

All procedures involving mice were carried out under the authority of personal licenses, in accordance with the Animal (Scientific Procedures) Act 1986. Severe Combined Immuno-deficient (SCID) mice were bred within the Biological Services Facility (BSF) of the University of Manchester. C57BL/6 mice were purchased from Envigo (Huntingdon, UK). C57BL/6 and SCID mice were housed and maintained in the Biological Services unit of the University of Manchester under Specific Pathogen Free conditions. A 12 hour light/dark cycle was maintained, with free access to food and water. All gavage and injection procedures were carried out in the morning between 9:00 and 11:00 to control for circadian effects.

### 5.2. Infection with T. muris

Eggs from the Edinburgh isolate of *T. muris* were obtained in the following manner: SCID mice were given 200 infective eggs by oral gavage. At 42 days post-infection (PI), mice were culled, and adult worms were extracted from caecum and proximal colon. Isolated worms were incubated for 24 hours in RPMI-1640 medium (Gibco, UK), in a 37°C incubator (with no atmospheric gas control). At 4 hours and 24 hours during incubation, eggs were removed from the media bathing worms, filtered in a 70 micron nylon strainer to remove debris and suspended in milliQ water. For experiments, animals were infected by oral gavage with 40 eggs of *T. muris* in a 200 µm bolus of milliQ water, and were kept under observation for the duration of infection.

### 5.3. MicroCT sample preparation

Infected mice were euthanised using CO_2_ in rising concentration. The caecum was dissected, and 0.5mm lengths of caecum tissue were fixed in 2.5 % glutaraldehyde, 4% PFA in 0.1M HEPES pH 7.2 for 24 hours. The samples were washed in ddH_2_O and were incubated in 1% aqueous Osmium tetroxide for one hour. After dehydration through graded ethanol, the samples were critical point dried.

For test samples stained with aqueous I_2_KI and Phosphotungstic Acid (PTA), we followed previously published methods [45]. Samples were stained in 1% aqueous solution of PTA, and aqueous 0.1% I_2_, 0.2% KI, and placed in a Eppendorf tube containing water to prevent drying and shrinkage during scanning.

### 5.4. MicroCT scanning

All microCT scans were executed on a Zeiss Versa 520 tomograph. An accelerating voltage of 50kV at 87.5mA and 4W was used. Different objective lenses (4X, 10X) were used depending on the desired final resolution. A detector binning of 2 was applied to the acquisition, and 1901 projections of 1.5 seconds each were collected. Filtered backprojection was used to reconstruct the tomogram.

### 5.5. Data analysis

Virtual slices and 3D volume renderings were generated and visualized in AVIZO 9.1. Watershed segmentation using AVIZO’s segmentation wizard gave a satisfactory initial rough segmentation which separated the worm, mucosa and submucosa. Manual annotation was used to generate kernels for the flooding algorithm. Due to local areas of low contrast especially in the head region, in almost all cases some manual alterations of the segmented worm were then required to provide the final segmentation included in figured.

For measurements of epithelial coverage, the total tunneled length of the stichosome, including epithelial breaks was estimated by use of AVIZO’s “centerline tree” algorithm on embedded portions of stichosome. Centerline tree extracted a non-branching spatial graph from the segmented worm which ran directly through the centre of the worm along the anterior-posterior axis. The proportion of the stichosome which was exposed by breaks in the epithelium overlaying the tunnel was calculated by manually separating the spatial graph of the embedded stichosome into either “exposed” or “unexposed” lengths which were subsequently measured. The cumulative length of the exposed stichosome was calculated and subtracted from the total tunneled length to obtain the proportion of the tunneled region of the stichosome which was exposed by epithelial breaks.

### 5.6. Histology

Mice were culled 35 days after initial infection and received an intraperitoneal injection (10ml/kg) of BrDU labelling reagent (no. 00-0103, Invitrogen, Camarillo CA) 40 minutes before culling. The caeca were fixed in methacarn, before transferring to 100% ethanol, paraffin embedding and sectioning (5 micron thickness slices). Worms with head down morphology were found by serial sectioning and searching of tissue sections, and stained both immunohistochemically for BrDU and using Haematoxylin and Eosin.

BrDU staining was carried out by the ABC method using the mouse on mouse kit (Vector:Peterborough UK) developed with the DAB system, using haematoxylin as a counter stain. A biotinylated primary antibody was used for direct detection without signal amplification (Biolegend 339809). Positive staining was confirmed by reference to the testing information provided by the manufacturer, as well as comparison with previously achieved results by our lab.

## 6. Acknowledgments

We acknowledge the Engineering and Physical Science Research Council (EPSRC) for funding the Henry Moseley X-ray Imaging Facility through grants (<u>EP/F007906/1</u>, <u>EP/F001452/1</u>, <u>EP/I02249X</u>, <u>EP/F028431/1</u>, <u>EP/M022498/1</u>, and platformgrant <u>EP/M010619/1</u>) within the Henry Royce Institute. PJW acknowledges support from the European Research Council grant No <u>695638 CORREL-CT</u>. KJE’s laboratory is supported by grants from both the Medical Research Council UK grant MR/NO22661/1 (https://mrc.ukri.org/) and Biotechnology and Biological Sciences Research Council grant BB/P018157/1 (https://bbsrc.ukri.org/). The authors wish to thank the staff in the FBMH EM Core Facility for their assistance and the Wellcome Trust for equipment grant support to the EM Facility. We further acknowledge Carl Zeiss AG for funding the studentship of JO.

